# Observing weak adaptation of duckweeds to their local microbiome depends on local pondwater

**DOI:** 10.1101/2025.05.12.653557

**Authors:** Ava Rose, Anna M. O’Brien

## Abstract

Populations can locally adapt to both the biotic and abiotic factors of an environment, but observing adaptation to biotic factors can sometimes depend on the abiotic conditions, and vice versa. One important aspect of the biotic environment is the microbiome: interactions between microbiomes and their hosts are critical for host fitness and trait expression. Hosts may adapt to their local microbiomes, and hosts may depend on interactions with microbes for adaptation to the local environment. Using *Lemna minor* (duckweed) as a model host organism, we examined differences in host fitness when grown in local and nonlocal microbiomes and in local and nonlocal water. We experimentally recombined duckweeds, microbes, and water from 4 different ponds around Durham, New Hampshire in well-plate microcosms in a growth chamber. Duckweed, microbe, and water source all affected microbial and duckweed growth, as well as duckweed traits. However, we observed only weak positive local adaptation that resulted in higher frond final area when duckweeds were paired with their local water and microbes. We also found that microbial growth was reduced when duckweeds were paired with microbes or water from their local site.

## INTRODUCTION

Populations often evolve in ways that increase their fitness in their local habitat, while experiencing a decay in fitness in other habitats due to drift or tradeoffs (Kawecki & Ebert, 2004; Hereford, 2009; Blanquart et al., 2013). Biotic factors can influence local adaptation, but whether they consistently do so has remained a debate. Meta-analyses have reached opposing conclusions: that leaving biotic interactions intact strengthens the signal of local adaptation (Briscoe-Runquist et al., 2020), and that biotic interactions do not alter signals of local adaptation (Hargreaves et al., 2020).

For macroorganisms, one important local biotic habitat may be the microbiome: the community of microorganisms that live on, inside, or near larger host organisms (Berg et al., 2020). Host-associated microbiomes alter fitness and trait expression in a wide variety of host species (e.g., Turnbaugh et al., 2008; Kaiko & Stappenbeck, 2014; Gould et al., 2018). For example, microbes often influence the timing of plant phenological events ranging from germination to fruiting across studied plants, regardless of microbe type, the nature of the interaction, and other factors (Lu et al., 2018; O’Brien et al., 2021). Indeed, the effects of microbiomes on plants are of particular interest, as we hope that engineering microbial treatments could increase crop yields and agricultural productivity (Qiu et al., 2019).

Some trait and fitness effects that microbiomes elicit in plants may derive from local adaptation. Plants may adapt to locally-associated microbes (Rúa et al., 2016), and reciprocally, microbes can adapt to their locally-associated plants (Batstone et al., 2020). However, local adaptation between plants and associated microbes or microbiomes can be conditional on the environment experienced, on the match of tested microbes to the full natural microbiome, and even on the environmental source of the plants and microbes (Johnson et al., 2010; Petipas et al., 2020; O’Brien & Laurich et al., 2024; O’Brien & Sawers et al., 2024).

The common pond plant *Lemna minor*, hereafter “duckweed,” has recently gained popularity as a model organism for host-microbiome interaction experiments (Ishizawa & Kuroda et al., 2020; Ishizawa & Tada et al., 2020; Anneberg et al., 2023; Jewell et al., 2023; Wei & Tan, 2023). Indeed, duckweed has been used as a model for plant biology since the 1950s (Acosta et al., 2021), largely because its rapid doubling time and small size make it easy to manipulate and ideal for experiments (Zhang et al., 2010), and its recent popularity has resulted in specialized experimental platforms and software (Mejbel & Simons, 2018; Cox Jr. et al., 2022; O’Brien et al., 2022; Kose et al., 2023; Subbaraman et al., 2024).

Here, we used *Lemna minor* and its associated microbiome as a model to quantify abiotic and biotic local adaptation by collecting duckweeds, water, and microbiomes from four sites and factorially recombining all materials in a “common garden” in the lab. We hypothesized that 1) duckweeds would have higher fitness when grown in both local biotic (microbiome) and local abiotic (water source) contexts than when grown in local contexts for only the biotic or abiotic factors (i.e. local microbiome but nonlocal water), or when grown in entirely nonlocal contexts (neither the microbiome nor the water from the home site). We also hypothesized 2) that inoculated microbiomes would reach higher cell densities when grown in local biotic and abiotic conditions (duckweeds and water from their home pond) than those grown in partially or entirely nonlocal contexts.

## MATERIALS AND METHODS

### Collection Sites, Samples, and Storage

Duckweeds, microbes, and water were collected from four locations across Durham, New Hampshire: Mill (M), Upper Mill (UM), Woodman (W), and the Durham Reservoir (DR, Table S1). Two pairs of sites were within the same watersheds: Mill and Upper Mill are in the Oyster River Watershed, and Durham Reservoir and Woodman are in the Pettee Brook Watershed.

Duckweed samples were collected from each of the four sites during the spring of 2022, and we used a single frond from each site to found a clonally propagated line in the lab. We expect these lines to be essentially isogenic, because duckweeds have low segregating diversity (Ho, 2018), and our lines were never observed to flower in our lab conditions. We created axenic cultures from these isogenic lines following an existing protocol (Laurich & Carlson, 2023). Briefly, duckweeds were agitated by vortexing for ∼5 minutes, placed in 1% bleach for 1 to 1.5 minutes, and then rinsed in sterile DI water three times. The first DI water rinse was 45 seconds long, and the second two were 5-10 minutes. The plants were transferred to sterile 0.5x Krazčič’s media until fronds grew. A few fronds were then placed in Krazčič’s media enriched with yeast and mannitol to check for failure of the procedure (microbial growth), and to rapidly amplify successful axenic duckweed cultures. Lines were grown in this media, in natural light conditions on a windowsill, refreshing media when growth slowed, until they were needed for experimentation.

Microbes were collected from each of the four sites between September and October 2023. Fresh gloves were rinsed with 70% ethanol and worn for the entire collection process. All tubes used for collection were sterile, and were rinsed 3 times in the source water prior to microbiome collection. Three 45 mL tubes of water with live local microbiomes were collected at each site. We centrifuged the collected water for 15 minutes at 4,000 rpm and 23℃. The supernatant was poured off, and the pellets were homogenized and resuspended in 2 mL of ∼25% sterile glycerol, then cryopreserved at -80℃ until needed. Before experimentation, inocula were resuscitated from the cryopreserved microbiomes by unfreezing and diluting each microbiome with 2 mL of sterile DI water. To create control treatments (sterile mock inocula with an identical glycerol concentration to live inocula), sterile 25% glycerol was diluted by half in sterile DI water.

Water samples were collected from each of the four sites from September to October 2023, at the same time as the microbiome samples. Three 40 mL samples of water were collected at each site. We placed the samples directly into a -20℃ freezer until the experiment set-up. Before experimentation, we removed excess dirt and debris using gravity filtration and paper filters. We then filter-sterilized water samples by passing them through a 0.2 µm filter.

### Experimental Design

To evaluate the impacts of local and nonlocal microbiomes and water on duckweed fitness, four axenic isogenic duckweed lines, four filter-sterilized water sources, and four revived microbiomes or a no microbiomes control were factorially recombined (80 treatments) in a randomized-block, microcosm experiment with 12 replicates in 10, flat-bottom 96-well plates. The randomized-block design simplified setup, which reduces errors and microbial cross-contamination. Each block contained one replicate of each of the 80 treatments, and contained 4 sub-blocks of 20 adjacent wells (4 rows by 5 columns), each randomly assigned to one duckweed line. Within each block, the field-collected water treatments were applied to random plate rows, and the cryo-preserved microbes or a no microbes treatment were applied to random plate columns (full experiment map is available in the GitHub repository).

We added each treatment to the plates following the randomized block design in a biosafety cabinet (REDISHIP Purifier Logic+ Class II A2, Labconco). We first added 205 μL of the sterilized field-collected water treatments, and then added single attached units of sterile duckweed fronds approximately the same in total surface area (ranging from 1-4 individual fronds). The plates were next sealed with Breath Easier seals (Diversified Biotech) to prevent cross-contamination when inoculating microbiome treatments. We restored microbiomes to their field-collected concentrations by adding 9.1 µL of microbial inocula (or of sterile mock inocula for the control, no microbes treatment) to the 205 µL of pond water already in the wells. The inocula treatments (live or mock) were added to each well by punching through the Breath Easier seal with a pipette tip. The plates were then sealed with BreathEasy seals (Diversified Biotech) and gently covered with the plate’s lid to slow evaporation in the growth chamber.

Once assembled, the experimental plates were placed in a Percival® model AR-75L3 growth chamber for 12 days. Lights and temperature cycled daily, with 16 hours at 350 µmoles/m^2^/s and 23℃, followed by 8 hours of darkness at 18℃. The plates were placed on a stage inside the growth chamber that was positioned over an imaging system (4 cameras connected to Raspberry Pis, as in Kose et al., 2023). Images of the plates were taken at the end of the experiment (Day 12), and analyzed with DuckPlate software to measure the surface area of duckweed fronds within each microcosm well (Kose et al., 2023). Frond area increases as the number of individuals increases and as individual quality (size) increases, and is therefore a commonly used fitness metric in duckweeds (O’Brien et al., 2020). After day 12, 200 µL of the remaining water was measured for optical density at 600 and 420 nm (OD_600_ and OD_420_) with a BioTek Cytation 5 plate reader. Optical density at 600 nm correlates with cell density in duckweed microbiomes (O’Brien et al., 2020).

### Data analysis

To test whether duckweed fitness was highest when duckweeds were paired with both water and microbes from their home site, and whether total microbial growth was highest when microbiomes were paired with both duckweeds and water from their home site, we used linear models. First, the 80 unique treatments were simplified into a matched treatment score based on whether, and if so how many, biological materials came from the same site, polarized by either the duckweed source or the microbe source. When polarizing by the duckweed source, the scores were based on whether the source site for the other materials was the same as the duckweed for: neither the water nor the microbiome (0), either the water or the microbiome (1), or both the water and the microbiome (2). When polarizing by the microbe source, the scores were based on whether the source site for the other materials was the same as the microbiome for: neither the water nor the duckweed (0), either the water or the duckweed (1), or both the water and the duckweed (2).

Statistical analysis was performed using linear models in R (R Core Team, 2021), with package MCMCglmm (Hadfield, 2010). Models were fit for the following equation for each response variable and each polarization (by duckweed or microbe source): *y* ∼ 𝛼 + 𝛽_*T*_*T* + *N*_*D*_(*0*) + *N*_*M*_(*0*) + *N*_*W*_(*0*), where 𝛼 is the intercept, T refers to the matched treatment score, and each *N*(*0*) parameter is a random effect for duckweed, microbe, or water source (D, M, W, respectively). Random effects for each treatment material’s source are included because strong effects of material source (e.g. high and low quality environmental matrix, maternal effects, etc.) can interfere with the correct interpretation of tests for local adaptation (Blanquart et al., 2013).

Various studies have found that sites within the same watershed share some commonality in their microbiomes, or other biotic communities (URycki et al., 2022; Comte et al., 2018). Because of this, the microbiomes or duckweed genotypes sampled from the same watershed may be more similar to each other than those from different watersheds. We therefore also fit models including a matched score for how many treatments matched the duckweed or microbe home watershed, again depending on which was used to polarize the match score. We then re-fit all models including this parameter: *y* ∼ 𝛼 + 𝛽_*T*_*T* + 𝛽_*T*W_ *T*_w_ + *N*_*D*_(*0*) + *N*_*M*_(*0*) + *N*_*W*_(*0*), where *T*_w_ refers to the watershed treatment score.

We further tested the main effect of inoculating with any microbe as opposed to uninoculated (no microbes) treatments for each measured response variable. To force microbially inoculated treatments to be treated the same by the model, we dropped the random effect of source microbes, using model: *y* ∼ 𝛼 + 𝛽_𝐼_𝐼 + *N*_*D*_(*0*) + *N*_*W*_(*0*), where *I* indicates the presence of any microbes inoculated onto duckweeds in microcosm wells. We also estimated main effects of duckweed, water, and microbe source singly (including uninoculated treatments), each in turn, leaving the other two treatment variables as random effects. E.g. for duckweed source site effects this model was *y* ∼ 𝛼 + 𝛽_*D*_*D* + *N*_*M*_(*0*) + *N*_*W*_(*0*). For all tested models, we report p-values to evaluate whether fitted effects are biologically significant (pMCMC values), and we report means and standard errors to visualize results.

If microbiomes shift the values of some response variables differently from others, they would also be expected to alter the correlations among response variables. Most of our measured response variables were weakly positively correlated to each other experiment-wide (rho = 0.16 to 0.36), except for the two optical density measures, which were very strongly positively correlated (rho = 0.96). For one optical density measure (600 nm), frond greenness, and frond area, we tested if pairwise correlations differed in microcosms filled with different water sources (*y* ∼ 𝛽_*W*_: 𝑥 + *N*_*D*_(*0*) + *N*_*M*_(*0*) + *N*_*W*_(*0*), for each unique 𝑥 and *y* pair of response variables). Similarly, we also tested if pairwise correlations between response variables differed when microcosms were inoculated with different microbiomes (𝛽_*M*_ instead of 𝛽_*W*_), or when different duckweed genotypes were added to microcosms (𝛽_𝑃_ instead of 𝛽_*W*_).

## RESULTS

Duckweed frond area, greenness, and both optical density measures varied substantially across the 80 unique treatments (Figures 1, S1). There were significant differences at p < 0.05 for one or more duckweed, water, or microbiome source for most response variables, except that greenness did not differ across duckweed or water source, and neither optical density measure differed across microbiome source (Figure S2). This highlighted the importance of including duckweed, water, and microbiome source site as random effects in other models. Compared to microcosms that were not inoculated with microbiomes, microcosms that were inoculated with microbiomes from any source had higher optical density (p < 0.05 for the natural log of optical density at both wavelengths, Figure S3). Uninoculated wells are not expected to have microbial cells, but contamination at the experiment setup stage does sometimes occur, as does measurement error, and both likely contributed to the few uninoculated treatments that showed higher optical density (Figure 1). Compared to microcosms that were not inoculated with microbiomes, microcosms that were inoculated with microbiomes from any source also had duckweeds with greater frond greenness (p < 0.05), and duckweeds that grew marginally faster than uninoculated duckweeds (p < 0.1, Figure S3).

**Figure 1:**
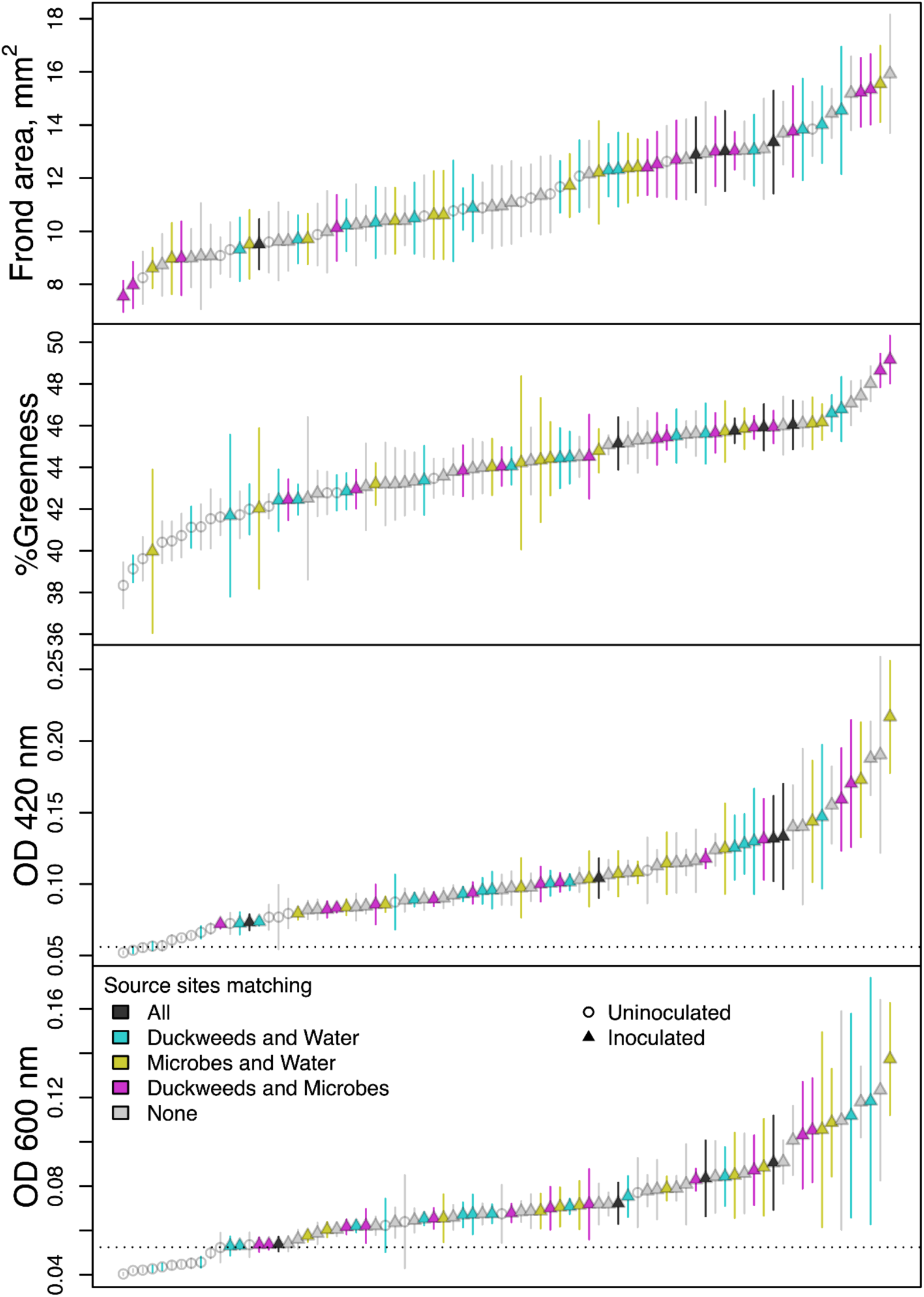
All treatment means for all response variables: duckweed frond area, duckweed frond percent green in pixel RGB values in images, and optical density at 420 nm and 600 nm in media. Points represent the average on day 14 of the experiment for each treatment category, while bars represent the standard error. N=12, except for optical density measures from three treatments, where N=11 due to NA values. Colors represent the type of match for source sites for the treatment (dark grey when all source sites matched, blue when only duckweed and water source matched, yellow when only microbe and water source matched, pink when only duckweed and water source matched, and light grey when no sources matched. Open circles are used for treatments in which no microbes were inoculated. The dashed line in the optical density panel is the average of blank media readings. See also Figure S1.

Only some of the variation across treatments was predicted by our tests for local adaptation. When treatments were increasingly drawn from the same source site as duckweeds (from 0, 1, or 2 of two additional treatments: microbes and water), microcosms had marginally more duckweed final area (p < 0.1), marginally less microbial growth (optical density at 600 nm, p < 0.1), marginally lower optical density at 420 nm (p < 0.1), and non-significantly greener duckweed fronds (Figure 2). When treatments were increasingly drawn from the same source site as the microbes (from 0, 1, or 2 of two additional treatments: duckweeds and water), no significant trends in the response variables were detected. However, microcosms trended towards more duckweed final area and greener duckweed fronds when all source sites were the same as the microbe source (all n.s., Figure 2, no trends for optical density). These observed, weak local adaptation effects suggest that local adaptation between duckweeds and microbes from their source site, and local adaptation of duckweeds to water from their source site together affect frond final area positively, and microbial growth negatively. Adding a numeric score for the number of source materials sharing a watershed did not substantially alter trends, though it did increase the significance of the frond area result, and decrease the significance of the optical density at 420 nm and microbial growth results (Figure S4, watersheds were Pettee Brook for Durham Reservoir and Woodman, and Oyster River for Upper Mill and Mill Pond).

**Figure 2:**
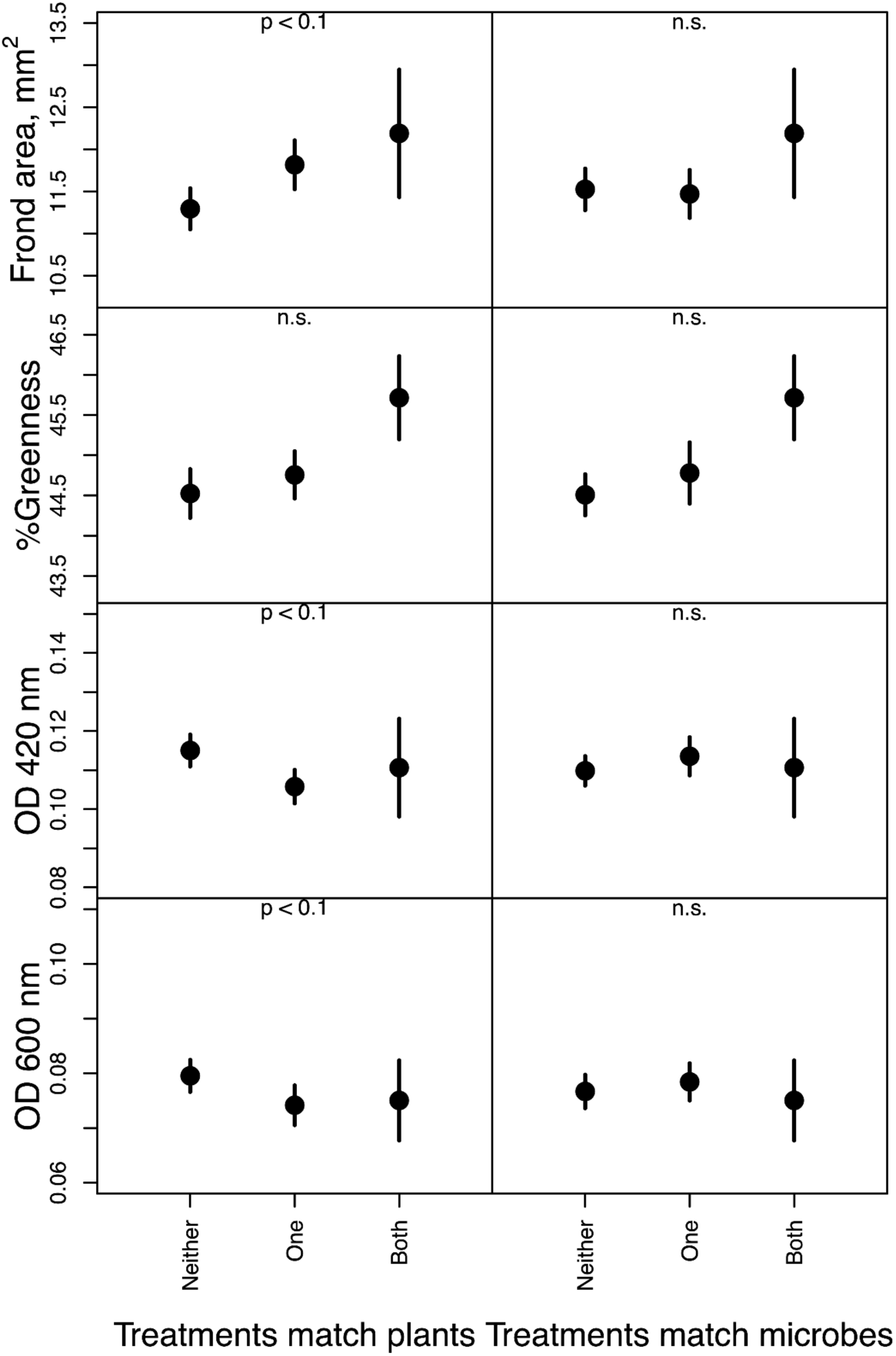
Duckweed frond area, duckweed frond percent green in pixel RGB values in images, and optical density at 420 nm and 600 nm in media. Points represent the average on day 14 of the experiment for each treatment category, while bars represent the standard error. Plots on the left show whether response variables changed as other treatments increasingly matched the plant source site, while plots on the right show whether response variables changed as other treatments increasingly matched the microbe source site. “*” indicates that the p-value of the increasing match (slope, 𝛽_*T*_) was different from 0 with p <0.05. Optical density measures are proxies for microbial cell density: optical density at 420 nm is expected to be increased by microbes with photosynthetic pigments and potentially fragmented host cells, while optical density at 600 nm is not. N = 432 for neither the water nor microbe source matches the plant source, and neither the water nor plant source matches the microbe source. N = 288 for either the water or microbe source matches the plant source, and either the water or plant source matches the microbe source. N = 48 for when all three sources match, and is the same data between the left and right columns.

Despite effects of duckweed source, microbiome source, and water source on most response measures, correlations among response variables were similar across treatments. However, some were significantly different from zero in only certain treatments. Positive correlations between frond greenness and frond area were always significant and did not differ across treatments (Figure 3). Positive correlations between frond area and the natural log of optical density at 600 nm did not differ from zero when the plants were from Durham Reservoir, the microbes were from Mill Pond, or the water was from Upper Mill. The positive correlation between frond greenness and the natural log of optical density at 600 nm was stronger when microcosms were inoculated with microbiomes from Durham Reservoir or Upper Mill than when microcosms were inoculated with microbiomes from Mill Pond, which instead yielded a non-significant negative correlation between greenness and microbial density (Figure 3). The correlation between frond greenness and the natural log of optical density at 600 nm was not significantly different from zero when the duckweeds were from Woodman, or when the water was from Woodman or Upper Mill.

**Figure 3:**
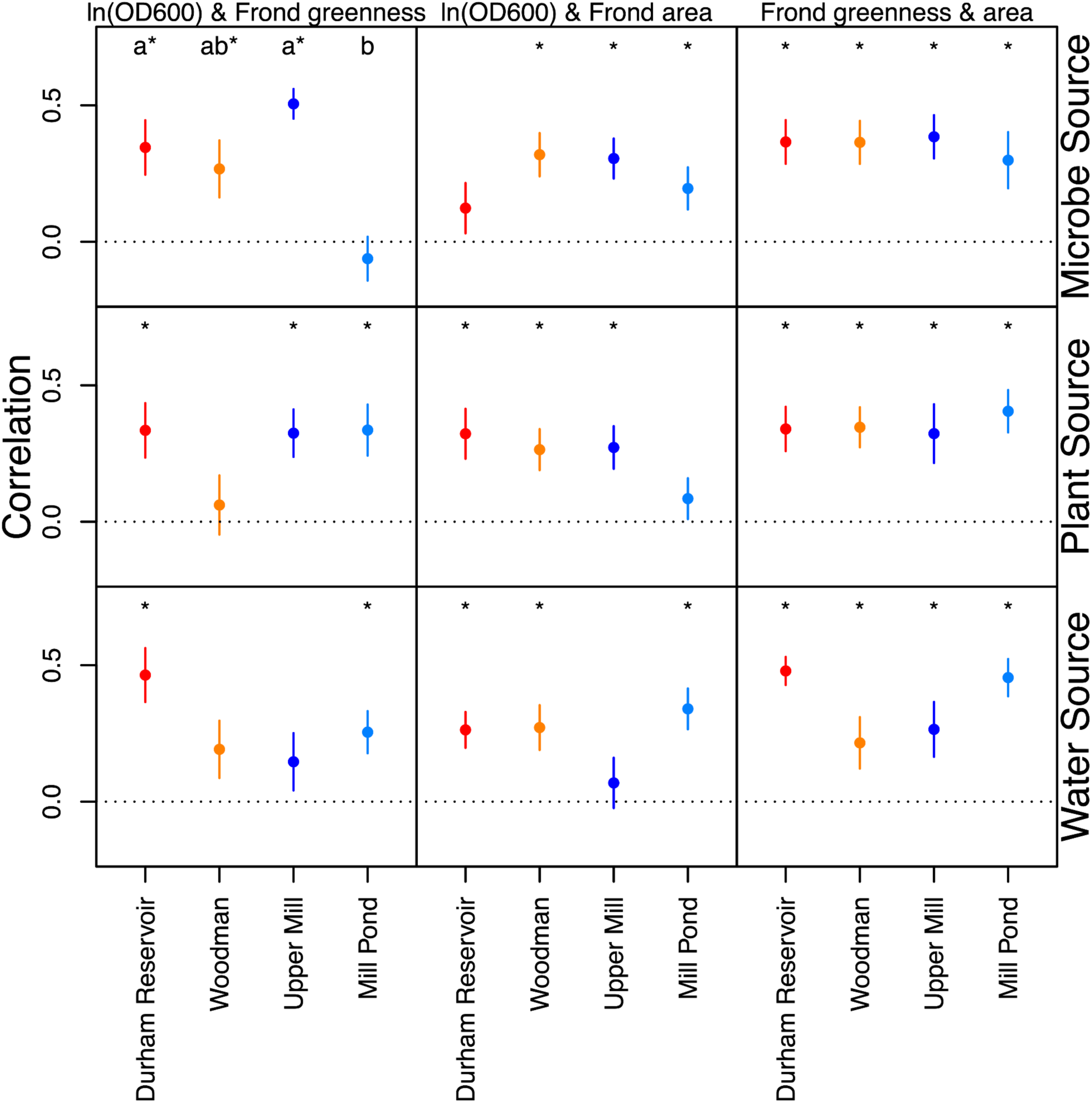
Correlations between three of the four measured response variables (the natural log of optical density at 600 nm, frond greenness, and frond area), averaged by source site for microbes, plants, and water separately. Correlations were calculated within each treatment (inoculated treatments only, 64 treatments, 12 datapoints each), then correlations were averaged for microbial, plant, and water treatments separately (points, bars are plus or minus one standard error of the mean, 16 treatments in each point). “*” indicates that mean was significantly different from zero in linear model results, and letters (when present) indicate significant differences in correlations between measured response variables across treatments.

## DISCUSSION

Local adaptation between species may result in higher fitness when individuals are paired with local versus non-local partners (Vos et al., 2009; Batstone et al., 2020). However, local adaptation can be conditional on the environment in which it evolved (Briscoe-Runquist et al., 2020). For example, we expect to see contingencies in local adaptation if adaptation to biotic interactions ameliorates stressors found only in the local environment (O’Brien et al., 2018; Petipas et al., 2021). Indeed, adaptation between plants and their local microbiome is sometimes only observed in the home environmental conditions (Johnson et al., 2010; Petipas et al., 2020; Iriart et al., 2024; Brady & Farrer, 2024). Or, interpreted reciprocally, part of plant local adaptation to the home environment can include local adaptation to microbes. Here, we found this to be the case in duckweeds. Duckweeds had weak, contingent signals of local adaptation to home site microbes and water: our metric of duckweed fitness was marginally higher when duckweeds were grown in both their home water and home microbiome, as compared to duckweeds grown with no local sources of microbes or water (Figure 2).

Previous work also found weak signals for local adaptation between duckweeds and their microbiomes, but had cultured microbiomes in the lab before testing, and had found that composition deviation between the cultured and field microbiomes appeared to erode the observable local adaptation (O’Brien & Laurich et al., 2024). We therefore expected to find a stronger signal of local adaptation in patterns of duckweed fitness when applying whole field microbiomes with limited manipulation, but we did not.

Aspects of the physical and biotic environment that define the home environment are often continuous. Sites that are nearer and further may be more ecologically similar, and indeed we often find that local adaptation is actually a quantitative phenomenon (Montalvo & Ellstrand, 2001; Wang et al., 2010; Wilczek et al., 2014). We therefore considered that the scale of local adaptation may be larger than a single pond, as there may be microbial or physicochemical similarities within the same watershed at low riverine distances (Payne et al., 2017; Liu et al., 2024). However, we also did not see a clear increase of local adaptation in both duckweed and microbial growth when widening the spatial definition of “home” (Figure S4).

Many other studies have found that individual microbe species sometimes locally adapt to their hosts or environments. For example, mycorrhizal fungi acquired more carbon from their local soil than non-local fungi (Johnson et al., 2010), *Saccharomyces paradoxus* matched to their climate conditions had higher fitness (Leducq et al., 2014), rhizobia matched with the host genotypes they evolved on induced more nodules (Batstone et al., 2020, Murray-Stoker & Johnson, 2024), and bacteriophages adapted to infect their local hosts in soil on a centimeters scale (Vos et al., 2009). We did not measure individual microbial species’ fitness, but we did measure total community growth. Total microbial growth was not higher when microbes were matched to their home host or home water environment. Indeed, the only marginally significant trend was that microbes grew more if the duckweeds in the microcosm did not come from the same site as the microbes or the water (Figure 2). Still, individual species abundances within total cell density may not reflect the total community results, and so we can only speculate about local adaptation or maladaptation of microbes to local duckweeds or local water.

Positive fitness feedbacks, where increased growth in one interactor results in increased growth or quality in the other, are expected to be critical drivers of evolution in positive species interactions (Foster & Wenseleers, 2006; Frederickson, 2013; Preussger et al., 2020). Total microbial growth was positively linked to duckweed fitness or frond greenness in most, but not all treatments in our experiment (Figure 3), suggesting that positive fitness feedbacks may be present in only some sites, conditional, or weak. If positive fitness feedbacks between duckweeds and their microbes are weak or variable, then this could limit the opportunities for local adaptation between duckweeds and microbes to provide fitness benefits, and explain the weak trends we observed here.

While we did not find a strong signal of local adaptation between duckweeds and microbes, we did find that the specific combinations of host genotype, microbiome, and water environment resulted in a wide variation across all response variables (Figures 1, S1). As for many other organisms, it is clear that manipulating the microbiome offers the opportunity to alter duckweed traits. Indeed, microbial manipulation remains an exciting avenue of product development for plants in general (Compant et al., 2025), and the bio-inoculants industry offers many products with microbes that are not local to the sites at which they will be used (Jack et al., 2021; Ladau et al., 2025). If local adaptation between plants and their microbiomes is generally weak, as for our duckweeds (Figure 2), there will be limited growth benefits of using microbes naturally associated with hosts. Yet, there can be unintended effects of non-local bioinoculant species, and they may escape into natural ecosystems (Mawarda et al., 2020; Jack et al., 2021). In general, even when local microbiomes are not best, they may also not be any worse than non-local microbiomes (Harrison et al., 2017; Brady & Farrer, 2024). Going forward, it may still be the safer ecological choice to bioprospect for probiotic strains in locally derived microbes.

## Acknowledgments

The authors wish to acknowledge funding support by the National Science Foundation BIO-MCB (Award #2300059) and United States Department of Agriculture Hatch project number NH 00724, collections at sites managed by the UNH Office of Woodlands and Natural Areas and the Town of Durham, and valuable discussion with other O’Brien Lab members (Alex Trott, Admas Berisso, Ciana Lazu, Alyssa Daigle, Ethan Morgan).

## Author contributions

AR and AMO contributed to the conception and design of the experiment. AR led the acquisition of data. AR provided the initial manuscript draft and data analysis, and all authors contributed to revising the draft and data analysis, and approved the final version to be published.

## Data Accessibility

Data and scripts are available at https://github.com/amob/Ladpt-Water-Microbe-MS-analysis

## SUPPLEMENTARY MATERIALS

**Table S1:** Geographic coordinates for sampled sites, and their respective watersheds.

| **Watershed** | **Site name** | **Latitude** | **Longitude** |
| --- | --- | --- | --- |
| Pettee Brook | Durham Reservoir | 43.1476 | -70.9441 |
| Pettee Brook | Woodman Road | 43.1374 | 70.9192 |
| Oyster River | Mill Pond | 43.1308 | -70.9215 |
| Oyster River | Upper Mill | 43.1225 | -70.9253 |

**Figure S1:**
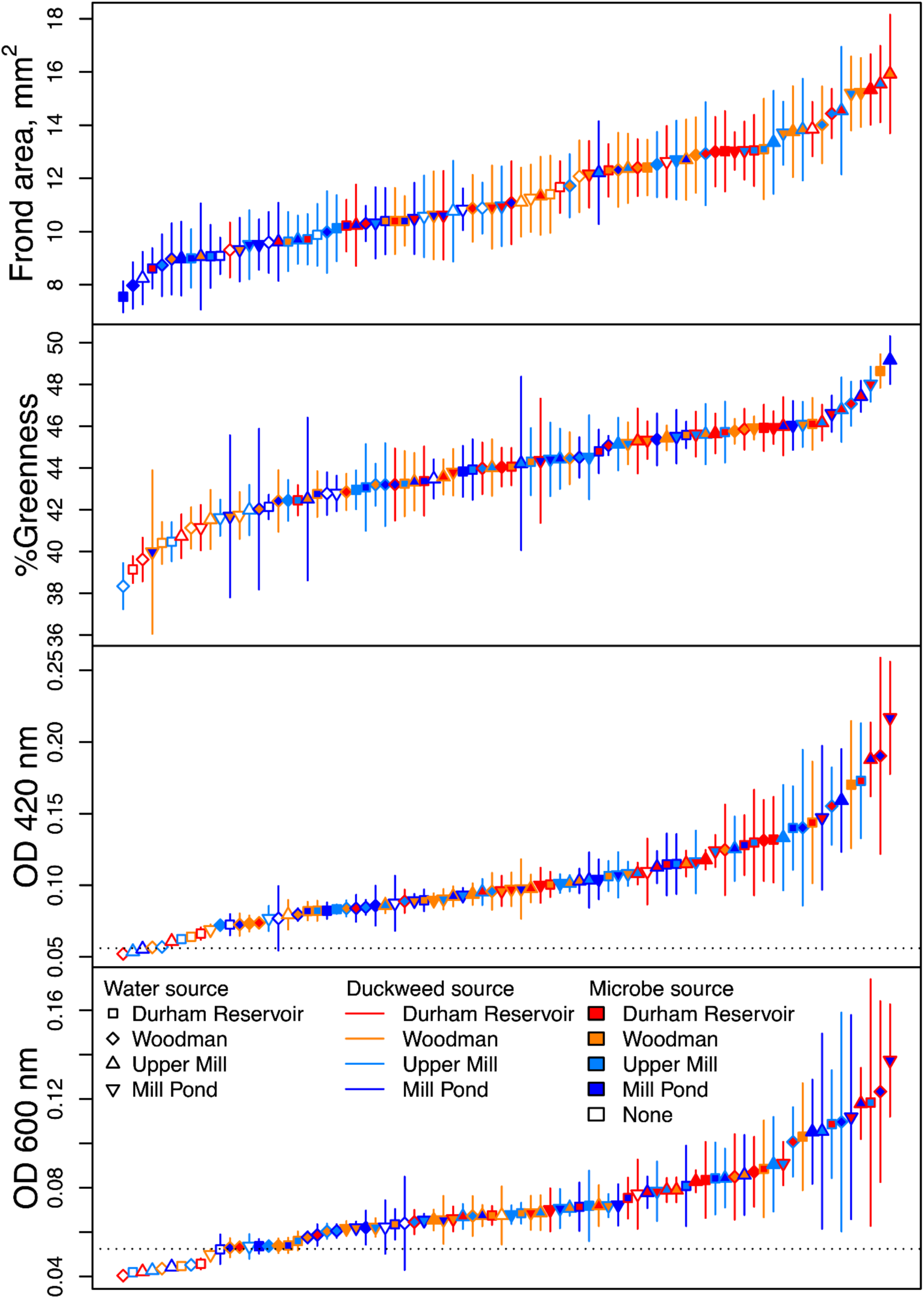
All treatment means for all response variables: duckweed frond area, duckweed frond percent green in pixel RGB values in images, and optical density at 420 nm and 600 nm in media. Points represent the average on day 14 of the experiment for each treatment category, while bars represent the standard error. N=12, except for optical density measures from three treatments, where N=11 due to NA values. Data are as in Figure 1, but treatments are indicated differently. Here, points are colored by source site for microbes, bars are colored by source site for duckweeds, and plotting symbols indicate the source site for the water. The dashed line in the optical density panel is the average of blank media readings.

**Figure S2:**
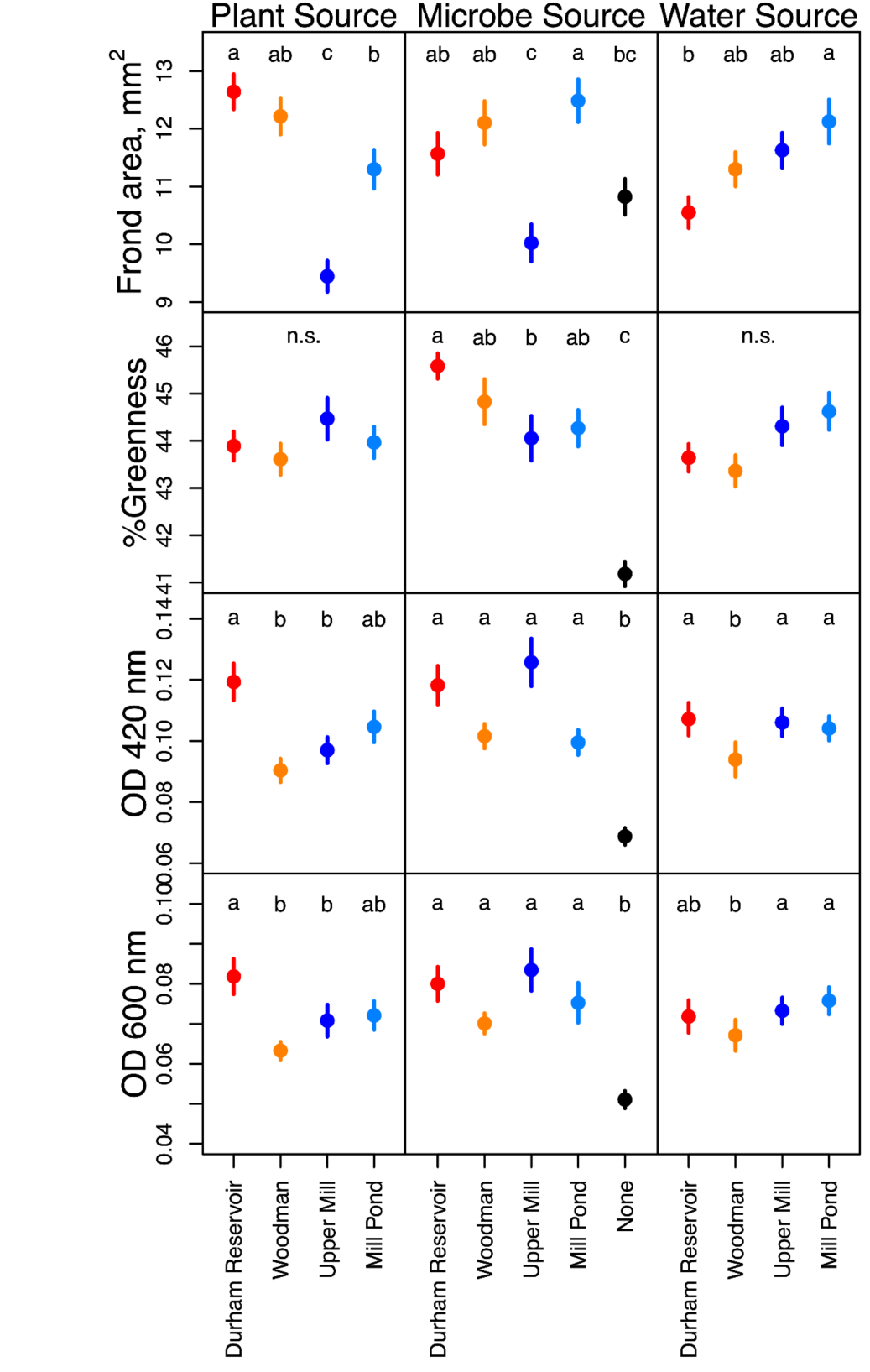
Means for each treatment, averaged across the others for all response variables: duckweed frond area, duckweed frond percent green in pixel RGB values in images, and optical density at 420 nm and 600 nm in media. Points represent the average on day 14 of the experiment for each treatment category, while bars represent the standard error. Each plant and water treatment has N=240; each microbe treatment has N = 192, except for optical density measures, where there were 3 NA values.

**Figure S3:**
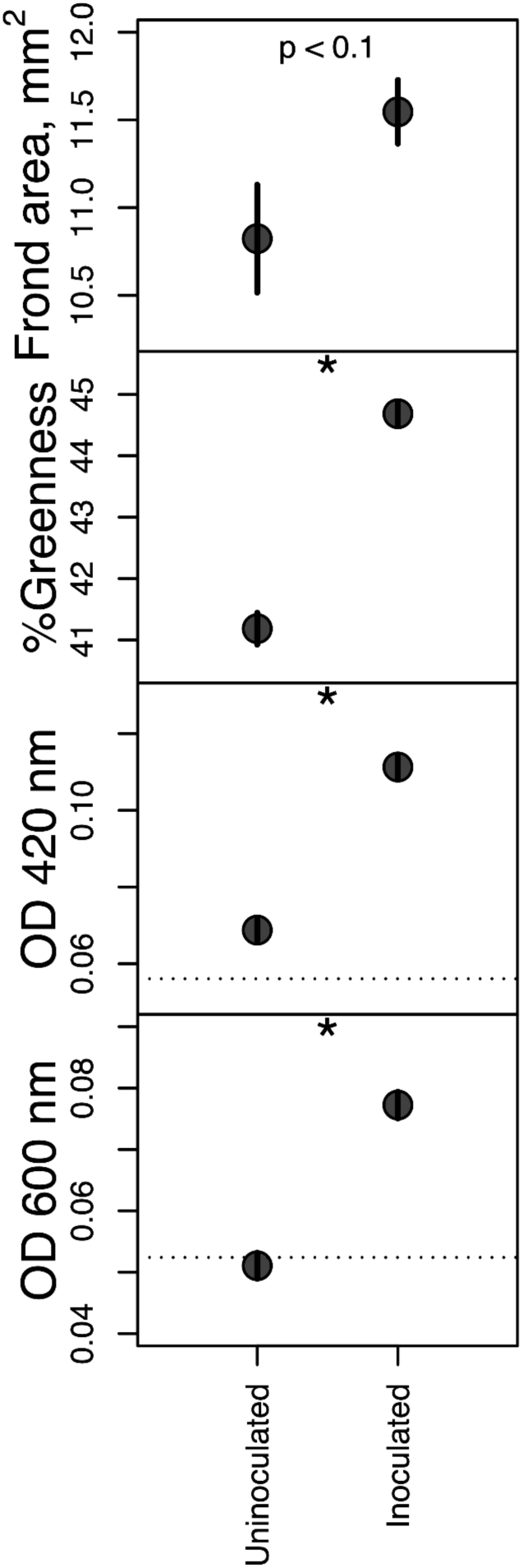
Duckweed frond area, duckweed frond percent green in pixel RGB values in images, and optical density at 420 nm and 600 nm in media, separated by whether microcosms were inoculated with microbes from any source, or received no microbes (uninoculated). Points represent the average on day 14 of the experiment for each treatment category, while bars represent the standard error. “*” indicates that the p-value of inoculation effect (slope, 𝛽_𝐼_) was different from 0 with p <0.05. Optical density measures are proxies for microbial cell density: optical density at 420 nm is expected to be increased by microbes with photosynthetic pigments, and/or fragmented duckweed cells, while optical density at 600 nm is not. N = 768 for inoculated microcosms, and N = 192 for uninoculated microcosms.

**Figure S4:**
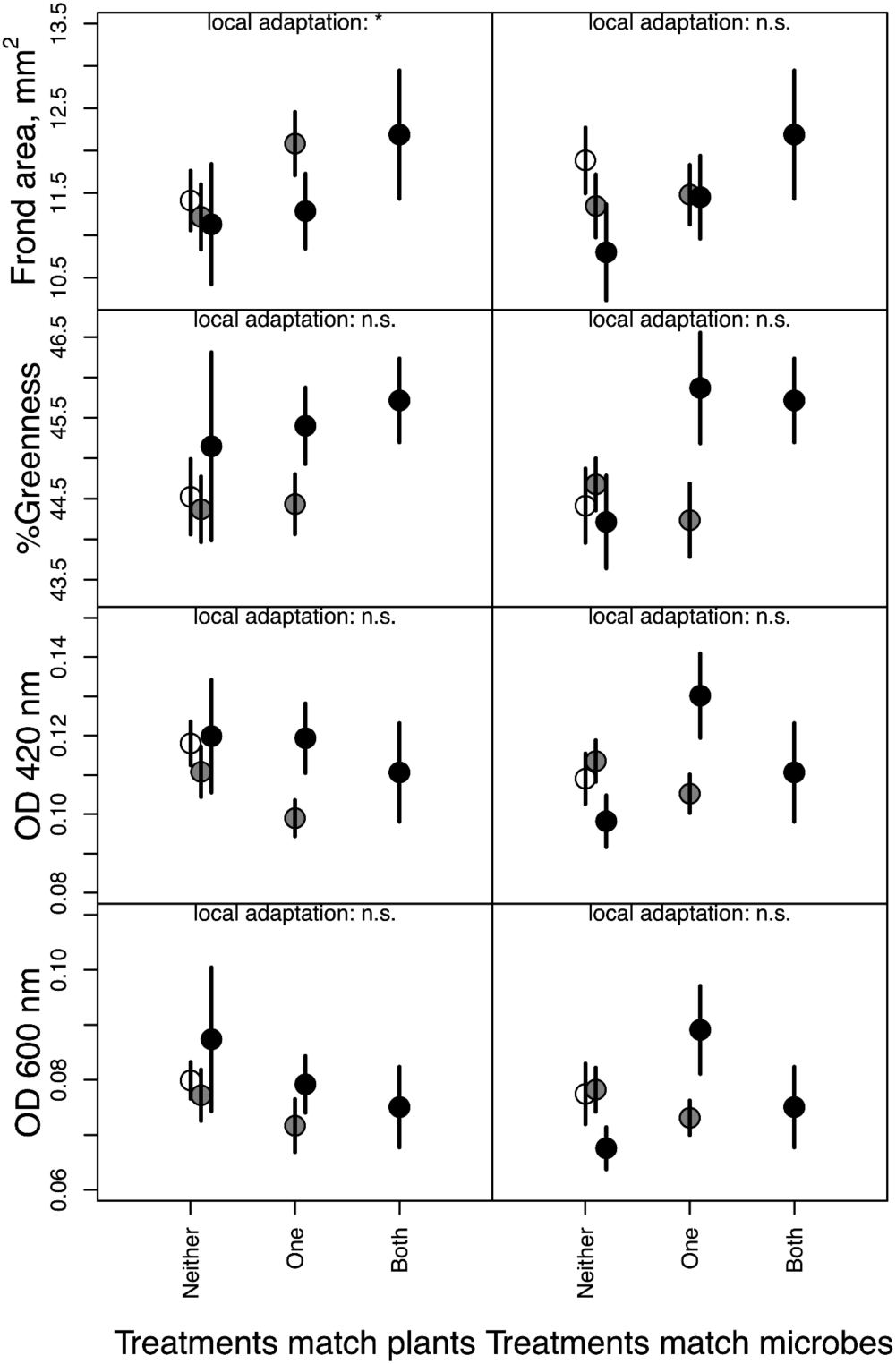
Data as in Figure 2, except that means and standard errors are further split by whether the watershed of the focal treatment (duckweeds on the left, microbes on the right) matches the watershed of neither of the other treatments (unfilled circles), one of the other treatments (grey circles), or all of the other treatments (filled circles). “*” indicates that the p-value of the increasing match (slope, 𝛽_*T*_) was different from 0 with p <0.05. Effects of watershed match were never significant (𝛽_*T*w_). N = 432 for neither of the other two sources matching the focal treatment’s source, and of these, 192 and 48 had one or both treatments’ source sites in the same watershed as the focal treatment’s source, respectively. N = 288 for when one of the other treatment’s source sites matches the focal treatment source, and of these, 96 had both of these sites from the same watershed as the focal treatment’s source. N = 48 for when all three sources match, and is the same data between the left and right columns, and in Figure 2.

## REFERENCES

Acosta, K., Appenroth, K. J., Borisjuk, L., Edelman, M., Heinig, U., Jansen, M. A., … & Lam, E. (2021). Return of the Lemnaceae: Duckweed as a model plant system in the genomics and postgenomics era. The Plant Cell, 33(10), 3207–3234.

Anneberg, T. J., Turcotte, M. M., & Ashman, T. L. (2023). Plant neopolyploidy and genetic background differentiate the microbiome of duckweed across a variety of natural freshwater sources. Molecular Ecology, 32(21), 5849–5863.

Batstone, R. T., O’Brien, A. M., Harrison, T. L., & Frederickson, M. E. (2020). Experimental evolution makes microbes more cooperative with their local host genotype. Science, 370(6515), 476–478.

Berg, G., Rybakova, D., Fischer, D., Cernava, T., Vergès, M. C. C., Charles, T., … & Schloter, M. (2020). Microbiome definition re-visited: old concepts and new challenges. Microbiome, 8, 1–22.

Blanquart, F., Kaltz, O., Nuismer, S. L., & Gandon, S. (2013). A practical guide to measuring local adaptation. Ecology Letters, 16(9), 1195–1205.

Brady, M. V., & Farrer, E. C. (2024). The soil microbiome affects patterns of local adaptation in an alpine plant under moisture stress. American Journal of Botany, 111(3), e16304.

Briscoe Runquist, R. D., Gorton, A. J., Yoder, J. B., Deacon, N. J., Grossman, J. J., Kothari, S., … & Moeller, D. A. (2020). Context dependence of local adaptation to abiotic and biotic environments: a quantitative and qualitative synthesis. The American Naturalist, 195(3), 412–431.

Compant, S., Cassan, F., Kostić, T., Johnson, L., Brader, G., Trognitz, F., & Sessitsch, A. (2025). Harnessing the plant microbiome for sustainable crop production. Nature Reviews Microbiology, 23(1), 9–23.

Comte, J., Culley, A. I., Lovejoy, C., & Vincent, W. F. (2018). Microbial connectivity and sorting in a High Arctic watershed. The ISME journal, 12(12), 2988–3000.

Cox Jr, K. L., Manchego, J., Meyers, B. C., Czymmek, K. J., & Harkess, A. (2022). Automated imaging of duckweed growth and development. Plant Direct, 6(9), e439.

Foster, K. R., & Wenseleers, T. (2006). A general model for the evolution of mutualisms. Journal of Evolutionary Biology, 19(4), 1283–1293.

Frederickson, M. E. (2013). Rethinking mutualism stability: cheaters and the evolution of sanctions. The Quarterly Review of Biology, 88(4), 269–295.

Gould, A. L., Zhang, V., Lamberti, L., Jones, E. W., Obadia, B., Korasidis, N., … & Ludington, W. B. (2018). Microbiome interactions shape host fitness. Proceedings of the National Academy of Sciences, 115(51), E11951–E11960.

Hadfield, J. D. (2010). MCMC methods for multi-response generalized linear mixed models: the MCMCglmm R package. Journal of Statistical Software, 33, 1–22. URL https://www.jstatsoft.org/v33/i02/.

Hargreaves, A. L., Germain, R. M., Bontrager, M., Persi, J., & Angert, A. L. (2020). Local adaptation to biotic interactions: a meta-analysis across latitudes. The American Naturalist, 195(3), 395–411.

Harrison, T. L., Wood, C. W., Borges, I. L., & Stinchcombe, J. R. (2017). No evidence for adaptation to local rhizobial mutualists in the legume Medicago lupulina. Ecology and Evolution, 7(12), 4367–4376.

Hereford, J. (2009). A quantitative survey of local adaptation and fitness trade-offs. The American Naturalist, 173(5), 579–588.

Ho, K. H. E. (2018). The effects of asexuality and selfing on genetic diversity, the efficacy of selection and species persistence. University of Toronto (Canada).

Iriart, V., Rarick, E. M., & Ashman, T. L. (2024). Rhizobial variation, more than plant variation, mediates plant symbiotic and fitness responses to herbicide stress. Ecology, 105(12), e4426.

Ishizawa, H., Tada, M., Kuroda, M., Inoue, D., Futamata, H., & Ike, M. (2020). Synthetic bacterial community of duckweed: a simple and stable system to study plant-microbe interactions. Microbes and Environments, 35(4), ME20112.

Ishizawa, H., Kuroda, M., Inoue, D., Morikawa, M., & Ike, M. (2020). Community dynamics of duckweed-associated bacteria upon inoculation of plant growth-promoting bacteria. FEMS Microbiology Ecology, 96(7), fiaa101.

Jack, C. N., Petipas, R. H., Cheeke, T. E., Rowland, J. L., & Friesen, M. L. (2021). Microbial inoculants: silver bullet or microbial Jurassic Park?. Trends in Microbiology, 29(4), 299–308.

Jewell, M. D., Van Moorsel, S. J., & Bell, G. (2023). Presence of microbiome decreases fitness and modifies phenotype in the aquatic plant Lemna minor. AoB Plants, 15(4), plad026.

Johnson, N. C., Wilson, G. W., Bowker, M. A., Wilson, J. A., & Miller, R. M. (2010). Resource limitation is a driver of local adaptation in mycorrhizal symbioses. Proceedings of the National Academy of Sciences, 107(5), 2093–2098.

Kaiko, G. E., & Stappenbeck, T. S. (2014). Host–microbe interactions shaping the gastrointestinal environment. Trends in Immunology, 35(11), 538–548.

Kawecki, T. J., & Ebert, D. (2004). Conceptual issues in local adaptation. Ecology Letters, 7(12), 1225–1241.

Kose, T., Lins, T. F., Wang, J., O’Brien, A. M., Sinton, D., & Frederickson, M. E. (2023). Accelerated high-throughput imaging and phenotyping system for small organisms. PLoS One, 18(7), e0287739.

Ladau, J., Fahimipour, A. K., Newcomer, M. E., Brown, J. B., Vora, G. J., Melby, M. K., & Maresca, J. A. (2025). Microbial inoculants and invasions: a call to action. Trends in Microbiology.

Laurich, J., & Carlson, C. (2023). Lemnaceae (Duckweed) culturing protocol. protocols.io

Leducq, J. B., Charron, G., Samani, P., Dubé, A. K., Sylvester, K., James, B., … & Landry, C. R. (2014). Local climatic adaptation in a widespread microorganism. Proceedings of the Royal Society B: Biological Sciences, 281(1777), 20132472.

Liu, X., Pan, B., Liu, X., He, H., Zhao, X., Huang, Z., & Li, M. (2024). Riverine microbial community assembly with watercourse distance–decay patterns in the north–south transitional zone of China. Journal of Hydrology, 628, 130603.

Lu, T., Ke, M., Lavoie, M., Jin, Y., Fan, X., Zhang, Z., … & Zhu, Y. G. (2018). Rhizosphere microorganisms can influence the timing of plant flowering. Microbiome, 6, 1–12.

Mawarda, P. C., Roux, X. L., Van Elsas, J. D., & Salles, J. F. (2020). Deliberate introduction of invisible invaders: A critical appraisal of the impact of microbial inoculants on soil microbial communities. Soil Biology & Biochemistry, 148, 107874.

Mejbel, H. S., & Simons, A. M. (2018). Aberrant clones: Birth order generates life history diversity in Greater Duckweed, Spirodela polyrhiza. Ecology and Evolution, 8(4), 2021–2031.

Montalvo, A. M., & N. C. Ellstrand. (2001). Nonlocal transplantation and outbreeding depression in the subshrub Lotus scoparius (Fabaceae). American Journal of Botany 88(2), 258–269.

Murray-Stoker, D., & Johnson, M. T. (2024). Mosaic of local adaptation between white clover and rhizobia along an urbanization gradient. Journal of Ecology, 112(5), 1150–1163.

O’Brien, A. M., Sawers, R. J., Ross-Ibarra, J., & Strauss, S. Y. (2018). Evolutionary responses to conditionality in species interactions across environmental gradients. The American Naturalist, 192(6), 715–730.

O’Brien, A. M., Laurich, J., Lash, E., & Frederickson, M. E. (2020). Mutualistic outcomes across plant populations, microbes, and environments in the duckweed Lemna minor. Microbial Ecology, 80(2), 384–397.

O’Brien, A. M., Ginnan, N. A., Rebolleda-Gómez, M., & Wagner, M. R. (2021). Microbial effects on plant phenology and fitness. American Journal of Botany, 108(10), 1824–1837.

O’Brien, A. M., Lins, T. F., Yang, Y., Frederickson, M. E., Sinton, D., & Rochman, C. M. (2022). Microplastics shift impacts of climate change on a plant-microbe mutualism: Temperature, CO2, and tire wear particles. Environmental Research, 203, 111727.

O’Brien, A. M., Laurich, J. R., & Frederickson, M. E. (2024). Evolutionary consequences of microbiomes for hosts: impacts on host fitness, traits, and heritability. Evolution, 78(2), 237–252.

O’Brien, A. M., Sawers, R. J., Gasca-Pineda, J., Baxter, I., Eguiarte, L. E., Ross-Ibarra, J., & Strauss, S. Y. (2024). Teosinte populations exhibit weak local adaptation to their rhizosphere biota despite strong effects of biota source on teosinte fitness and traits. Evolution, 78(12), 1991–2005.

Payne, J. T., Millar, J. J., Jackson, C. R., & Ochs, C. A. (2017). Patterns of variation in diversity of the Mississippi river microbiome over 1,300 kilometers. PloS One, 12(3), e0174890.

Petipas, R. H., Wruck, A. C., & Geber, M. A. (2020). Microbe-mediated local adaptation to limestone barrens is context dependent. Ecology, 101(8), e03092.

Petipas, R. H., Geber, M. A., & Lau, J. A. (2021). Microbe-mediated adaptation in plants. Ecology letters, 24(7), 1302–1317.

Preussger, D., Giri, S., Muhsal, L. K., Ona, L., & Kost, C. (2020). Reciprocal fitness feedbacks promote the evolution of mutualistic cooperation. Current Biology, 30(18), 3580–3590.

Qiu, Z., Egidi, E., Liu, H., Kaur, S., & Singh, B. K. (2019). New frontiers in agriculture productivity: optimized microbial inoculants and in situ microbiome engineering. Biotechnology Advances, 37(6), 107371.

R Core Team (2021). R: A language and environment for statistical computing. R Foundation for Statistical Computing, Vienna, Austria. URL https://www.R-project.org/.

Rúa, M. A., Antoninka, A., Antunes, P. M., Chaudhary, V. B., Gehring, C., Lamit, L. J., …& Hoeksema, J. D. (2016). Home-field advantage? Evidence of local adaptation among plants, soil, and arbuscular mycorrhizal fungi through meta-analysis. BMC Evolutionary Biology, 16, 1–15.

Subbaraman, B., de Lange, O., Ferguson, S., & Peek, N. (2024). The Duckbot: A system for automated imaging and manipulation of duckweed. PLoS One, 19(1), e0296717.

Turnbaugh, P. J., Bäckhed, F., Fulton, L., & Gordon, J. I. (2008). Diet-induced obesity is linked to marked but reversible alterations in the mouse distal gut microbiome. Cell Host & Microbe, 3(4), 213–223.

URycki, D. R., Bassiouni, M., Good, S. P., Crump, B. C., & Li, B. (2022). The streamwater microbiome encodes hydrologic data across scales. Science of The Total Environment, 849, 157911.

Vos, M., Birkett, P. J., Birch, E., Griffiths, R. I., & Buckling, A. (2009). Local adaptation of bacteriophages to their bacterial hosts in soil. Science, 325(5942), 833–833.

Wang, T., O’Neill, G. A., & Aitken, S. N. (2010). Integrating environmental and genetic effects to predict responses of tree populations to climate. Ecological Applications, 20(1), 153–163.

Wei, N., & Tan, J. (2023). Environment and host genetics influence the biogeography of plant microbiome structure. Microbial Ecology, 86(4), 2858–2868.

Wilczek, A. M., Cooper, M. D., Korves, T. M., & Schmitt, J. (2014). Lagging adaptation to warming climate in Arabidopsis thaliana. Proceedings of the National Academy of Sciences, 111(22), 7906–7913.

Zhang, Y., Hu, Y., Yang, B., Ma, F., Lu, P., Li, L., … & Chen, S. (2010). Duckweed (Lemna minor) as a model plant system for the study of human microbial pathogenesis. PLoS One, 5(10), e13527.

